# A thrombus is formed by a gradient of platelet activation and procoagulant endothelium

**DOI:** 10.1101/2023.08.19.550692

**Authors:** Estelle Carminita, Julie Tourn, Lydie Crescence, Nicolas Brouilly, Glenn Merrill-Skoloff, Alexandra Mazharian, Christophe Dubois, Laurence Panicot-Dubois

## Abstract

**Introduction:** The contribution of platelets in thrombosis within microcirculation has been extensively documented in the literature. We previously showed, *in vivo,* that platelet activation revealed by intracellular calcium mobilization was a crucial step in the growth of thrombi following laser-induced injury, a model of thromboinflammation.

**Aim:** We employed a multimodal, correlative microscopy approach and computational biology to investigate the extent of platelet activation and the spatial distribution of platelets throughout a growing thrombus.

**Results:** We observed a reversible intracellular platelet calcium mobilization that correlates with the time a platelet resides during thrombus growth. Our bioinformatics analysis displayed three distinct platelet subpopulations resident within a thrombus: **(1)** resting, **(2)** partially activated, and **(3)** “fully” activated platelets. The spatial distribution of the platelet subpopulations in the thrombus creates a double gradient in both the transversal and longitudinal axis, with the maximal percentage of fully activated platelets close to the site of injury. However, these activated platelets did not express negative phospholipids. The injured endothelium was identified to play a vital role in activating the blood coagulation cascade in this model of thrombosis.

**Conclusion:** Following a laser-induced injury, thrombi are formed by a gradient of activated platelets from the injury site to the periphery of the thrombus. These different activation states of platelets throughout the thrombi regulate the biomechanics of the thrombus. The injured endothelium, rather than platelets, was identified to play a key role in the activation of the blood coagulation cascade in this model of thromboinflammation.

**Essentials:**

- Computational biology was used to analyze thrombosis.
- Non-activated, low- and fully-activated platelets are part of a thrombus.
- The activation of the platelets forms a gradient from the site of injury to the periphery.
- The endothelium, and not platelets, expressed negative phospholipids.

**Graphical abstract:** A thrombus is formed by a gradient of platelet activation and procoagulant endothelium

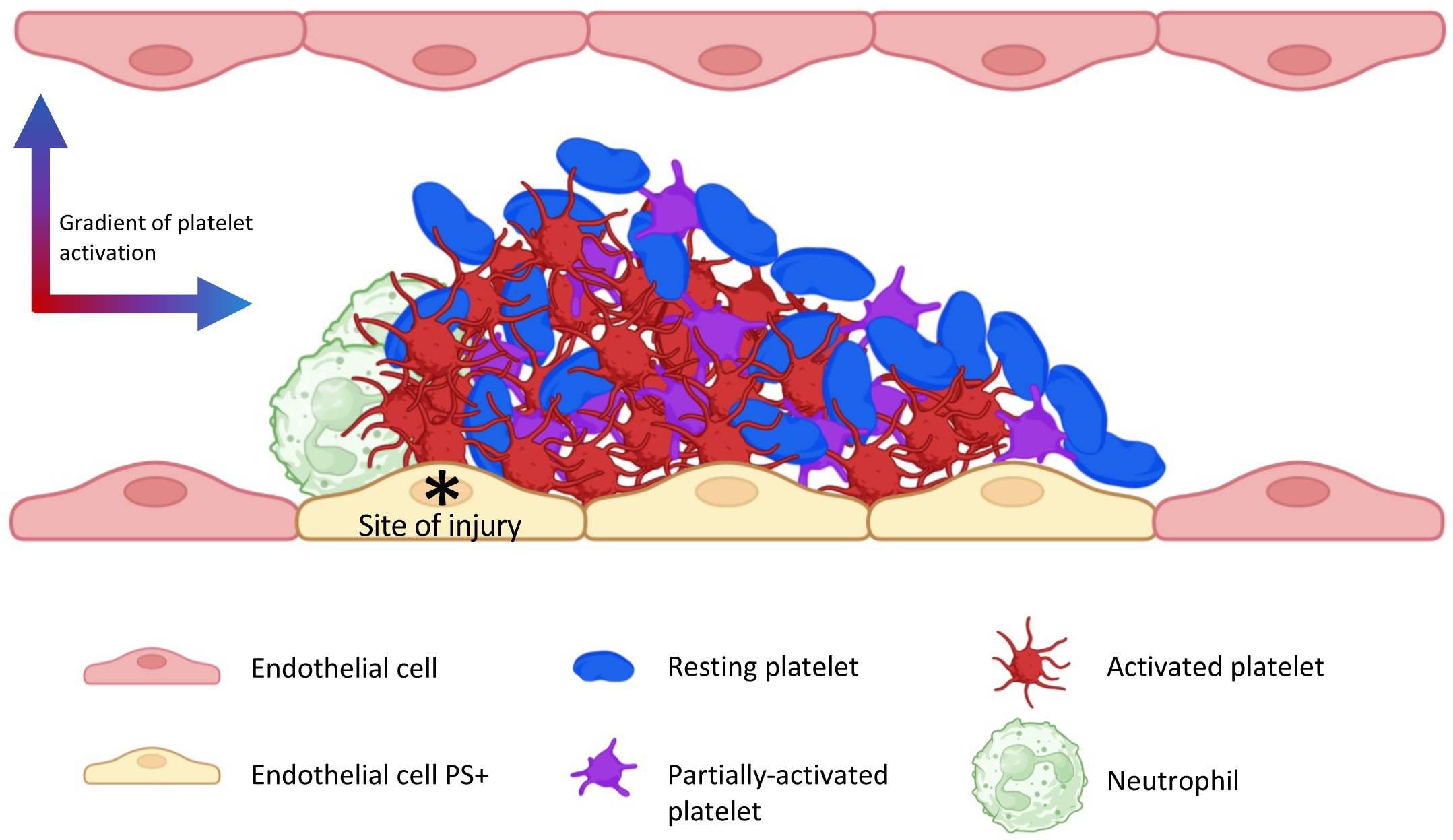

## Introduction

Adhesion and aggregation of platelets to an injured vessel wall are critical steps in thrombus formation. Once platelets become adherent, they are activated and recruit additional circulating platelets by secreting secondary agonists such as adenosine diphosphate (ADP) and thromboxane A2 (TxA2)[1][2] Upon activation, platelets change their shape: from an initially discoid structure, platelets can spread out by forming filopodia and lamellipodia[3]. The degree of this shape change depends on the degree of activation of the platelet: a weakly activated platelet will remain in its discoid form and is typically named a “primed platelet”. The procoagulant activity of platelets is one of the consequences of platelet activation and is characterized by platelet granule secretion and aggregation [4]. Procoagulant platelets such as "COATed” platelets (COllagen And Thrombin activated) are characterized by a high level of intracellular calcium, a loss of mitochondrial potential, the secretion and expression of proteins typically localized within granules now on the outer membrane surface as well as P-Selectin, the expression of negative phospholipids at their surface [5].

Platelet activation within a thrombus is, however, not uniform. The first evidence demonstrating this point came from *in vitro* studies using electron microscopy [6][7]. Within the last decade, it has become evident that platelets are not a homogeneous cell population but rather a heterogeneous assortment of subpopulations. These subpopulations differ in their activation, shape, and thus their functions [8]. This heterogeneity in platelet populations has prompted scientists to study the composition of platelet clots resulting from different experimental thrombosis models using *in vivo* mouse models in conjunction with *ex vivo* human sample preparations.

The study of the hierarchical structure of thrombi and the distribution of differentially activated platelet subpopulations is highly controversial and differs depending on the model used. Brass *et al*. [9] were the first to characterize two distinctive parts with a growing thrombus using the “Furie model” (a dye laser on the cremaster microcirculation of a living mouse): **(1)** An inner core close to the site of injury, composed of closely packed P-selectin positive platelets. This region’s primary agonist is thrombin, which results in fibrin deposition[10]. The core is hard to infiltrate with targeting antibodies [11]. The presence of procoagulant platelets in this core is unknown; however, our computational biology approach allows us to observe and quantify the platelet composition within the core of thrombi. **(2)** The second region defined by Brass was an outer shell formed by loosely packed platelets that do not express P-selectin. The primary platelet agonists in this region are ADP and TxA2 [12].

We have previously demonstrated that, when using the Furie model, activated neutrophils on endothelium act as the initiator of the Tissue Factor-dependent (TF) coagulation cascade needed for the accumulation and activation of platelets at the site of injury. Our results also emphasized the contribution of the adenosine triphosphate (ATP)/Adenosine ratio in the activation of both neutrophils and platelets [13][14], rendering the role played by the different platelet agonists within the core and shell of a thrombus even more complex than previously thought. In this manuscript, we studied the degree of activation of platelets throughout thrombi using multimodal, correlative microscopy and computational biology approaches with real-time laser scanning intravital confocal microscopy (LSCIM) and serial block-face scanning electron microscopy (SBF-SEM) in the Furie model of thromboinflammation. Based on our previous results, we hypothesize that, in this model, a thrombus could be formed by resting and activated but not procoagulant platelets.

We demonstrate that firm adhesion of platelets to developing thrombi requires calcium mobilization and show that rather than two separate sections (core and shell), a radial gradient of platelet activation is formed throughout the thrombus emanating from the thrombus core. We also showed using P2Y12 deficient mice that P2Y12 signaling is required for efficient platelet-platelet interaction, but its contribution to coagulation is limited. Indeed, fibrin generation is independent of the activation of platelets, but it is colocalized with the endothelium.

## Methods

### Mice

Wild-Type (WT) C57Bl/6JRj mice (5 to 9 weeks old) were obtained from Elevage Janvier (Le Genest-Saint-Isle, France). P2Y12-deficient mice (EM:02301, B6;129-P2ry12^tm1Dgen^/H) were purchased from DeltaGen Inc (San Mateo, CA, USA; EMMA, EM;02301). All animal care and experimental procedures were performed as recommended by the European Community Guidelines (directive 2010/63/UE) and approved by the Marseille Ethical Committee #14 (protocol number: APAFIS#20334-2019041811535225 and APAFIS#15334-2018060115491816V2).

### Antibodies and reagents

Antibodies and reagents were purchased from the following vendors. Anti-mouse GPIb antibody (Cat# X649, Emfret, Eibelstadt, Germany). Anti-mouse CD31 antibody (clone MEC13.3, BioLegend, London, United Kingdom). Isolectin GS-IB4 from Griffonia simplicifolia (Cat#I21412, ThermoFisher Scientific, Dreieich, Germany). Fura-2 AM and BAPTA-AM (Cat# F1221, Molecular Probes, Eugene, OR, USA and CAS# 126150-97-8, Calbiochem San Diego, CA, USA, respectively. Anti-mouse GPIbα (CD42b) used for platelet depletion (Cat# R300, Emfret, Eibelstadt, Germany). Annexin-V PE-Cy5 (cat# 1015, BioVision, Waltham, MA, USA).

### Preparation of washed mouse platelets and use *in vivo*

Washed platelets were prepared as previously described [13]. Three WT C57BL/6 mice were anesthetized (100 mg/kg ketamine, 25 mg/kg xylazine, and 0,25mg/kg atropine), and blood was collected in citrated buffer (1/9e) in the presence of 0,5mM prostacyclin and 0,02U/mL apyrase. Platelets were isolated by centrifugation at 450g for 15 minutes at 37°C and washed twice in Tyrode’s buffer in the presence of 0,04U/mL apyrase and 500nM PGI2 (cat#, manufacturer, city, country). Washed platelets were resuspended at 1x10^8^ platelets/mL in Tyrode’s buffer containing 0,2% BSA. Platelets were incubated with 3mM Fura-2 AM in the absence or presence of 50mM BAPTA-AM for 40 minutes in the dark at 37°C, centrifuged, and finally resuspended in Tyrode’s buffer at 1x10^8^ platelets/mL before injection into a recipient mouse. Resting platelets are visualized at a wavelength of 363nm and depicted in green. Activated platelets undergoing calcium flux are visualized at 335nm and depicted in red.

### Preparation of human platelet-rich plasma (PRP), aggregation test and scanning electron microscopy (SEM)

Human PRP and platelets were prepared as previously described [13]. Briefly, blood was centrifuged, and PRP was prepared in the presence of CaCl_2_ (2mM) and MgCl_2_ (1mM) for 30 minutes at 37°C. PRP was pre-incubated at 37°C in a cuvette using an aggregometer (APACT 4004, Ahrensburg, Germany). The aggregation assay was started with a 0% aggregation baseline. Then, PRP was exposed to ADP (12μM) or vehicle control. An aggregation test was performed for 5 minutes, followed by fixation with 4% PFA-Phosphate Buffer 0,1M pH 7,4 for one hour. Adhered resting or ADP-activated human platelets on coverslips treated with poly-lysine were coated with 20nm gold in 24-well flat-bottom plates. After dehydration, coverslips were coated with 20nm gold a second time and analyzed in a scanning electron microscope (Quanta 200, FEI, Eindhoven, Netherlands).

### Intravital microscopy and laser-induced injury

Intravital video-microscopy of the cremaster muscle microcirculation was done as previously described [13]. WT mice CR57BL/6 were pre-anesthetized with intraperitoneal ketamine (100mg/kg), xylazine (12,5mg/kg), and atropine (0,25mg/kg). A tracheal tube was inserted, and the mouse was maintained at 37°C. To maintain anesthesia, thiopental (12,5mg/kg every 30min) was administered through a cannula placed in the jugular vein. Following the incision of the scrotum, the testicle and cremaster muscle were exteriorized, and the cremaster was perfused with a thermo-controlled buffer (37°C). Micro-vessel data were obtained using an Olympus AX microscope with a 60X magnification water immersion objective. Digital images (640x480 pixels) were captured with a Cooke Sensecam CCD camera, followed by image analysis using SlideBook 6 (Intelligent Imaging Innovations, Denver, USA). Exogenous Fura-2 AM labeled platelets for the calcium experiments [13] or labeled antibodies for the intravital confocal experiments [14] were infused into the circulation of an anesthetized mouse. A vessel wall injury within the cremaster was induced with a dye-based pulsed nitrogen laser (MicroPoint, Photonics Instruments). Typically, 1 or 2 pulses were required to cause vessel injury and thrombus formation, as reported in the literature [13][14][15]. Multiple thrombi were studied in a single mouse for 1 hour with new thrombi formed upstream of earlier thrombi to avoid any contribution from thrombi generated earlier in the animal.

### Serial block-face scanning electron microscopy (SBF-SEM) and platelet morphology analysis

SBF-SEM sample preparation was performed as previously described [16]. Briefly, platelet thrombi were fixed using glutaraldehyde, ferrocyanide-reduced osmium tetroxide for post-fixation, thiocarbohydrazide and osmium tetroxide, followed by uranyl acetate and lead aspartate. The samples were dehydrated in a graded series of ethanol to absolute ethanol and acetone and embedded in Durcupan resin. Ultra-thin sections (90 nm) were obtained with an ultra-microtome for morphological examination using a transmission electron microscope (Tecnai G2, Thermo Fisher Scientific). Cremaster muscles with thrombi were mounted on an SBF pin, and the acquisition was performed upstream of the blood flow. The samples were visualized with an SEM microscope equipped with a VolumeScope SBF module (Teneo VS, ThermoFisher Scientific). The acquisition was performed at a thickness of 100nm and a pixel size of 10 nm. For the morphology analysis, we systematically observed and counted ∼1000 platelets at the site of injury in the thrombi. Each platelet was assigned a numerical identifier ‘1-1000’. The dataset was randomized, and ∼100 platelets were chosen to be segmented. Once segmented, the sphericity value of each platelet was calculated according to the following formula: S = 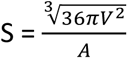 (V: Volume; A: Area) and platelet distribution depending on the activation state was established.

### Statistical analysis

Data are representative of at least three experiments for all the *in vivo* experiments. For the intravital experiments, the significance of the fibrin generation was demonstrated using a one-way ANOVA with Mann-Whitney test at 95% CI (Graphpad software). Regarding the colocalization statistics, an unpaired Student t-test was performed. For the electron microscopy experiments, statistics were performed with one-way ANOVA with Tukey’s multiple comparisons test at 95% CI (Graphpad software). ** P< 0.01.

## Results

### Characteristics of the platelet calcium mobilization *in vivo*

We previously described the intracellular calcium mobilization into platelets participating in a growing thrombus following a laser-induced injury [16]. To study the calcium flow, 250x10^6^ Fura-2 AM loaded platelets were infused into a recipient mouse [13][14][15][16]. To determine the characteristics of the calcium mobilization in single-bound platelets *in vivo,* we also performed experiments with lower ratios of labeled platelets to unlabeled platelets by injecting 1x10^6^ Fura-2 AM loaded platelets into mice. Under these conditions, we could follow individual platelets’ accumulation and calcium mobilization into a growing thrombus (**Figure 1A**). Only one spike of calcium was observed in single platelets. In the absence of calcium mobilization, platelets adhere but do not bind tightly enough in shear conditions (shear rate = ∼1500 s^-1^). Of the total number of labeled platelets detaching from the developing thrombus, 72% showed no evidence of calcium mobilization (**Suppl Video 1**).

**Figure 1:**
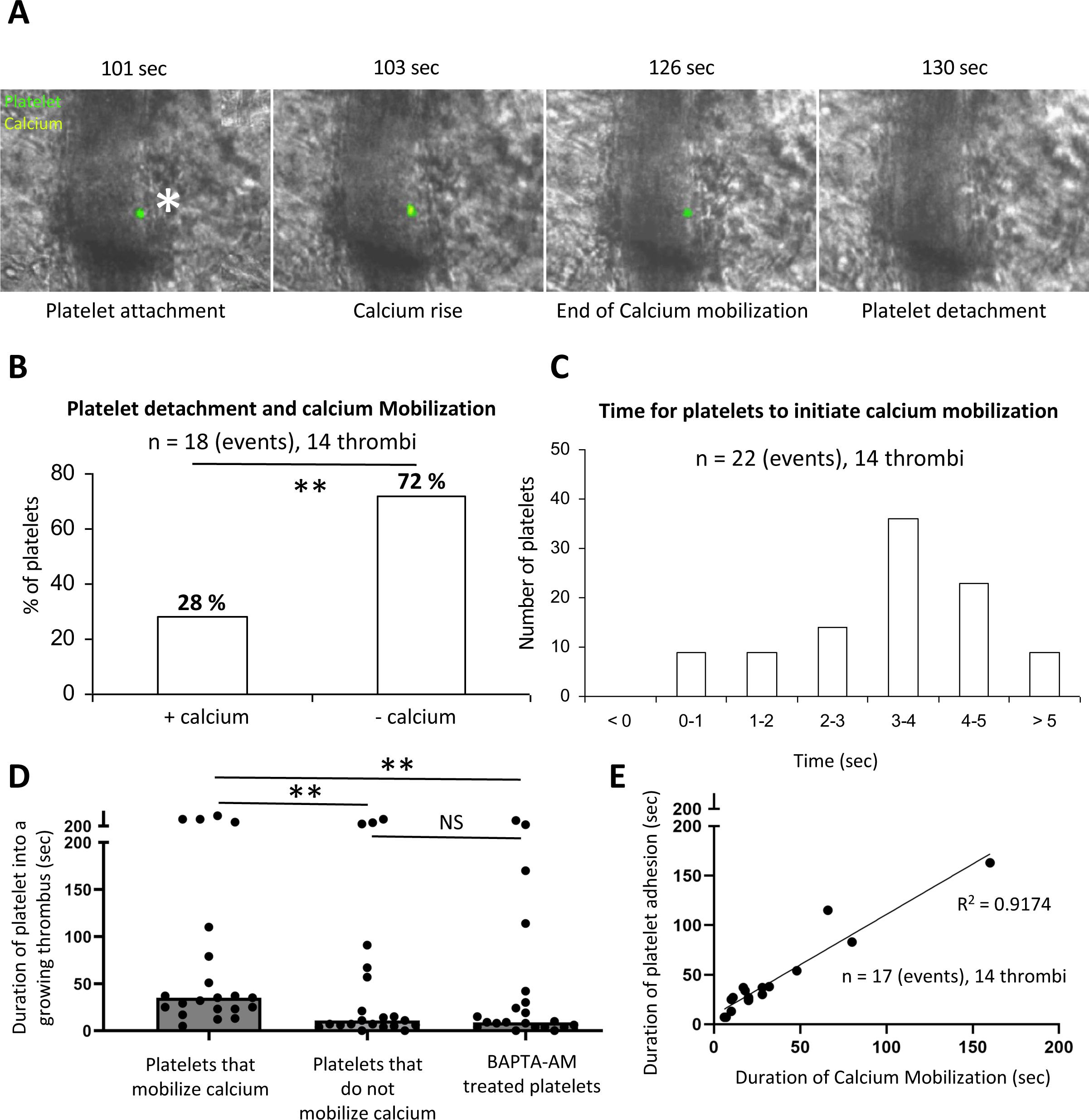
Intra-platelet calcium mobilization is responsible for the time-dependent incorporation of platelets in thrombi. (A): Representative images of the course of platelet calcium flux over time. Wild-type platelets were purified, labeled with Fura-2 AM, and reinjected into a recipient mouse through the jugular vein (1% of the total platelets were injected). Platelets are labeled in green, and the calcium flow inside the platelets is depicted in yellow. the width of the artery was typically 20 μm. (B): Quantitative analysis of the relationship between the calcium flux mobilization in platelets and their detachment from the thrombus (n=18 events, 14 thrombi, three mice). (C): Quantitative analysis of the time for platelets to mobilize calcium in the thrombus (n=22 events, 14 thrombi, three mice). (D): Graph represents the duration of the platelets inside the thrombus according to their calcium mobilization state (n=17 events, 14 thrombi, 3 mice). Data are shown as medians, and statistics were performed using a one-way ANOVA with Mann-Whitney test at 95% CI (E): Graph represents the correlation between the duration of platelets in the growing thrombus depending on the duration of the calcium mobilization (n=17 events, 14 thrombi, 3 mice). ** P< 0.01.

Nonetheless, some platelets (28%) that had undergone calcium mobilization detached from the thrombus and returned to circulation **(Figure 1B)**. Most of these platelets were associated with platelet emboli, including unlabeled platelets and platelets that had undergone calcium mobilization. The time interval from adhesion to calcium mobilization for each platelet varied from 1.0 to 12 sec, with a median of 3.5 sec **(Figure 1C)**, suggesting that a platelet could reversibly incorporate into a growing thrombus without any calcium mobilization. However, more than 90% of platelets that underwent calcium mobilization did so within 5 sec of adhesion. In vivo, a mean duration of 20 sec for calcium mobilization was calculated from multiple observations (n = 17), ranging from 7 to 165 sec (**Figure 1E**). Platelets that continued to mobilize calcium remain incorporated within the growing thrombus for a greater period (median of 35 sec) than platelets that did not mobilize calcium (median of 11 sec) or platelets treated with BAPTA-AM to inhibit calcium mobilization (median of 9 sec) **(Figure 1D)**. However, it remains likely that specific signaling processes initiated by high levels of intracellular calcium remain active even after calcium concentrations return to resting levels. The return to resting levels of calcium in the platelet resulted in decreased attachment from thrombi. There was a close correlation between calcium mobilization in an individual platelet and the duration associated with the developing thrombus, R^2^ = 0.9174 **(Figure 1E).**

To determine if platelets get differentially activated depending on their spatial orientation within the thrombus, we next performed LSCIM of Fura-2 AM loaded platelets accumulating at the site of injury (**Figure 2A**). We observed calcium mobilization in platelets present in the traditional “core and shell” of the thrombus (**Figure 2A**). To better analyze the activation of platelets according to their spatial orientation in the thrombus, we divided it into three different regions: **(1)** The first region was defined as close to the site of laser-induced injury, representing around 25% of the thrombus surface and corresponds to the core part of the thrombus. **(2)** The second region is the intermediate zone. **(3)** The third region, representing 50% of the thrombus surface, is at the periphery of the injury site (**Figure 2B**).

**Figure 2:**
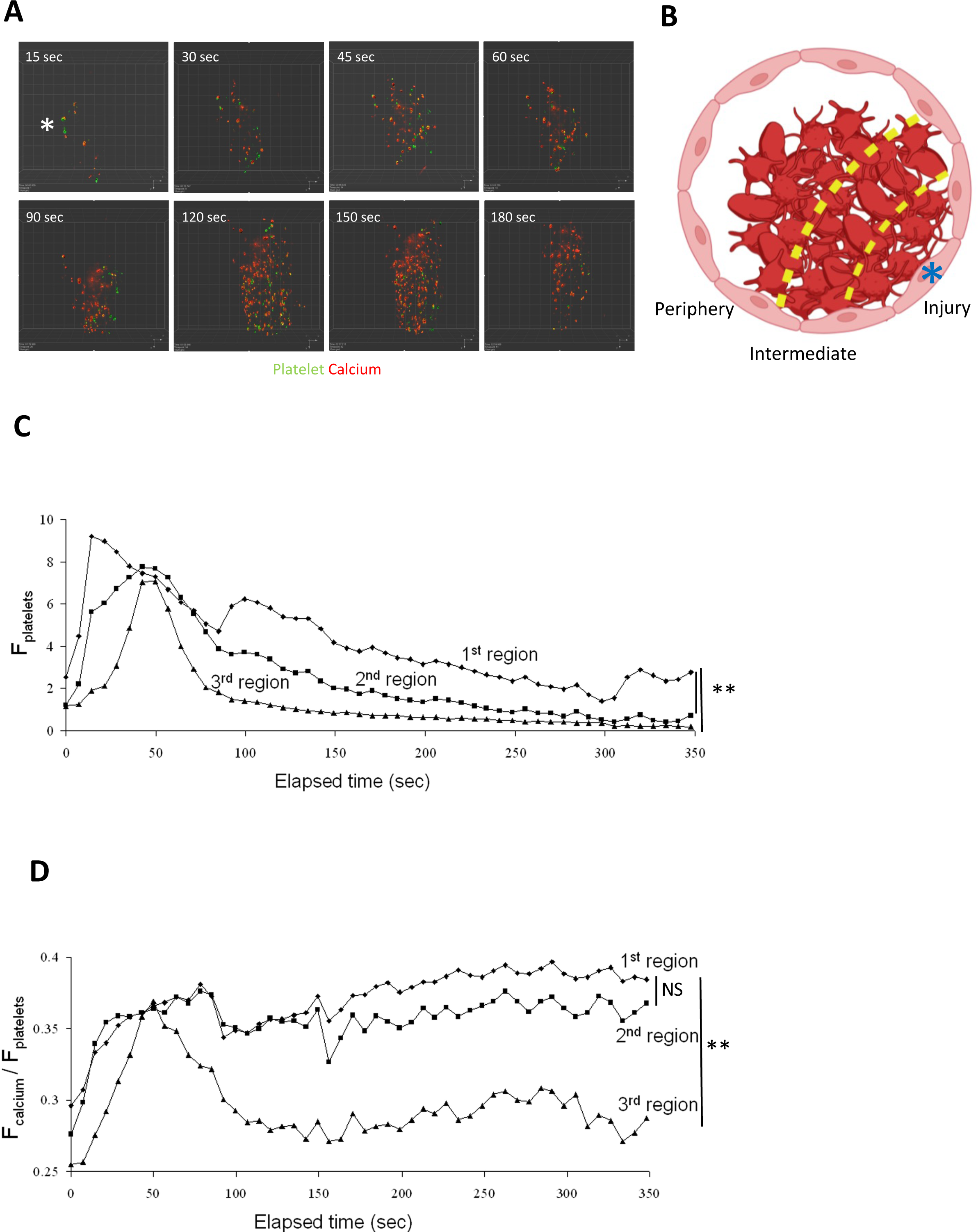
Platelets’ spatial organization within thrombi depends on calcium flux. (A): Representative images of the course of platelet calcium flux over time in LSCIM. Wild-type platelets were purified, labeled with Fura-2 AM, and reinjected into a recipient mouse through the jugular vein. Injected platelets represent 20% of the total platelets (representative of 18 thrombi observed in 3 different mice). Platelets are depicted in green and platelets mobilizing calcium are in red. The site of laser injury is indicated by an asterisk. (B): Representative schematic of thrombi and analyzed zones. Thrombi were divided into three zones: the first region is close to the injury, the second region is intermediate, and the third region is defined as the thrombus periphery. The site of laser injury is indicated by an asterisk. (C, D): Kinetics of platelet accumulation (C) and the ratio of calcium flow per platelet (D) over time for each previously described region. Dots represent the median of fluorescent signals (n=8). Data are shown as medians, and statistics were performed using a one-way ANOVA with Mann-Whitney test at 95% CI. ** P< 0.01.

When comparing the accumulation of platelets over time in the different regions, we observed that in the first region, only some platelets -stayed attached up to 350 seconds post-injury. On the contrary, close to 90% of the platelets in the third region were detached at 100 seconds post-injury (**Figure 2C**). These results correlate with the activation status of the platelets in the different regions. Whereas platelets still mobilized calcium in the first two regions, calcium flow into platelets was rapidly reversed in the third region and remained low after 100 seconds post-injury (**Figure 2D**). These results indicate that the intensity of calcium mobilization into platelets, which corresponds to the degree of platelet activation, may control the duration of platelet attachment in a growing thrombus. Activated platelets are present in the different parts of the thrombus.

### A radial gradient of platelet activation is observed by computational biology within a thrombus

We next performed, first *in vitro*, a control SEM experiment using human platelet-rich plasma (PRP) to observe the change of platelet morphology during an aggregation test (**Supp Figure 1A**). Under basal conditions (**Supp Figure 1A, top panel**), platelets remain quiescent in their resting state and are discoid. Following ADP stimulation (12μM), platelets progressively change their morphology, become partially activated with few filopodia (**Supp Figure 1A, middle panel**), and then form additional filopodia (fully activated) (**Supp Figure 1A, bottom panel**). Using segmentation and bioinformatic analysis, these results identify three different shapes and three activation states of platelets within the same platelet sample.

To investigate this further, we investigated the activation of platelets participating in a growing thrombus *in vivo* using volume scanning electron microscopy by serial block-face (SBF-SEM). This technique, coupled with segmentation and bioinformatic analysis, allowed us to study the cellular morphology of platelets within the thrombus. Briefly, we systematically annotated 1000 platelets at the site of injury in the thrombi. Each platelet was assigned a numerical identifier ‘1-1000’. The dataset was randomized, and more than one hundred platelets were segmented (**Supp Figure 1B**). As expected, discoid (solid arrows) and filopodia-presenting (dotted arrows) platelets were present in a thrombus (**Figure 3A**). We were able to distinguish by computational biology the three subpopulations of platelets detected *in vitro*: inactivated or "resting" platelets that remained discoid (**Figure 3B left panel**), activated platelets with several filopodia (**Figure 3B right panel**), and intermediate or “partially activated” platelets with a morphology that we categorize between fully activated and resting platelets (**Figure 3B middle panel**). The “partially activated” platelets display a lower number of filopodia compared to the fully activated platelets and are not discoid such as resting platelets. Next, we calculated the sphericity values S of each segmented platelet according to the formula: S = 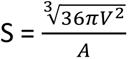 (V: Volume; A: Area). This geometric parameter varies between 0 and 1, with the value of 1 corresponding to a sphere. The calculation of the sphericity values of the platelets confirms our previous observations. Three subpopulations of platelets could be defined (**Figure 3C**): resting platelets with sphericity values between 0.86 and 0.57 (48/115 = 42%), partially activated platelets between 0.56 and 0.54 (15/115 = 13%), and fully activated platelets with a value between 0.53 and 0.33 (52/115 = 45%). We next investigated the presence of the three subpopulations of platelets in the three previously defined regions of the thrombus (**Figure 3C-D**). The "Injury" first region, containing the injured endothelium, comprised almost 87% of activated platelets. Of note, this region also contains a minority of partially activated and resting platelets (about 7% of each category). The intermediate region comprises 67% fully activated platelets, 10% partially activated platelets, and 23% resting platelets. Lastly, in the periphery region, most of the platelets were not activated (57%). However, we also observed partially and fully activated platelets in this region (**Figure 3E**). Taken together, these results indicate the existence of a transverse gradient of activation from the wound-injured endothelium-to the periphery of the vessel, with most of the activated platelets being found closest to the site of injury with resting platelets at the periphery of the thrombi.

**Figure 3:**
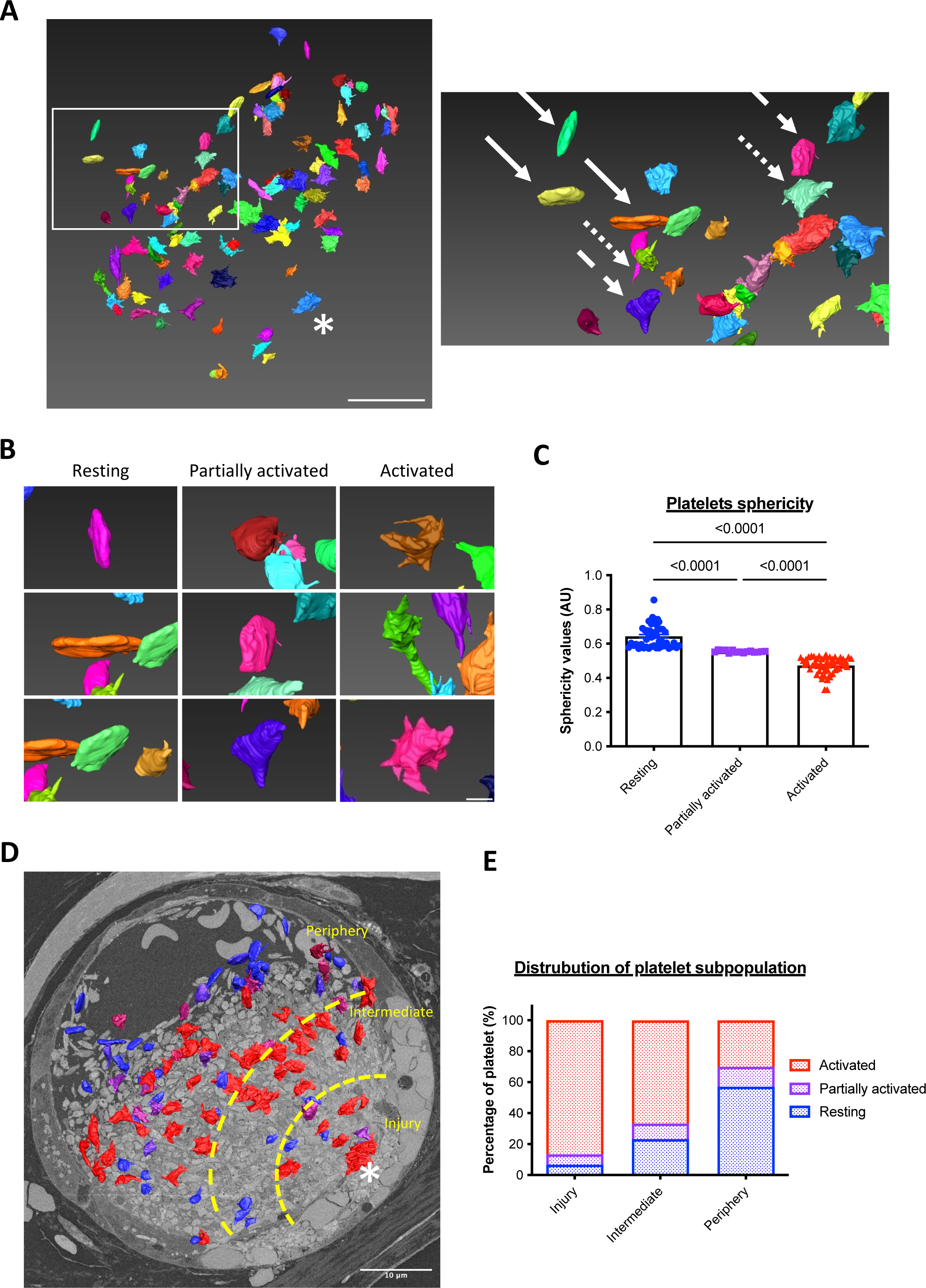
Platelets are distributed in the thrombus according to their activation state. (A): Representative image of the segmented 10% platelets within a thrombus following a laser-induced injury in SBF-SEM (left panel) (n=3). The site of laser injury is indicated by an asterisk. Scale bar: 10μm. Right panel: enlargement revealing three subpopulations of platelets differentially activated: resting platelets (solid arrows), partially activated (dashed arrows), and fully activated (dotted arrows). (B): Enlargements of the representative SBF-SEM image of the three subpopulations of platelets composing the thrombus. Three examples for each sub-population are shown schematically. Scale bar: 1μm. (C): Quantitative analysis of sphericity values of segmented platelets in the thrombus after a laser-induced injury. Each point corresponds to one sphericity value and, therefore, to an individual platelet, resting platelets (blue), partially activated (purple), and fully activated (red). One-hundred platelets have been analyzed. The mean +/- SEM is shown. Significance was determined using one-way ANOVA with Tukey’s multiple comparisons test. (D): Representative image of a thrombus following a laser-induced injury in SBF-SEM revealing the three parts studied: injury, intermediate, and periphery. Scale bar: 10μm. (E): Quantitative analysis of the distribution of platelets according to their state of activation. One thousand platelets were segmented and over a hundred platelets have been analyzed using the Amira software (Thermo Fisher Scientific).

We next studied the activation state of platelets in the longitudinal direction (**Figure 4A**). We continued to find the three previously described regions: the site of injury “Injury” (**Figure 4B left panel**), the middle region "Middle" (**Figure 4B middle panel**), and the tail of the thrombus “Tail” (**Figure 4B right panel**). The thrombus compactness values (**Figure 4C**) and the average distance between two neighboring platelets (**Figure 4D**) were calculated. Briefly, compactness is a geometric parameter calculated by a ratio between the area of the studied object and the total area of the field in which this object is located. Its value varies between 0 and 1, with 1 corresponding to a compact object with few empty spaces. The distance between platelets was also calculated.

**Figure 4:**
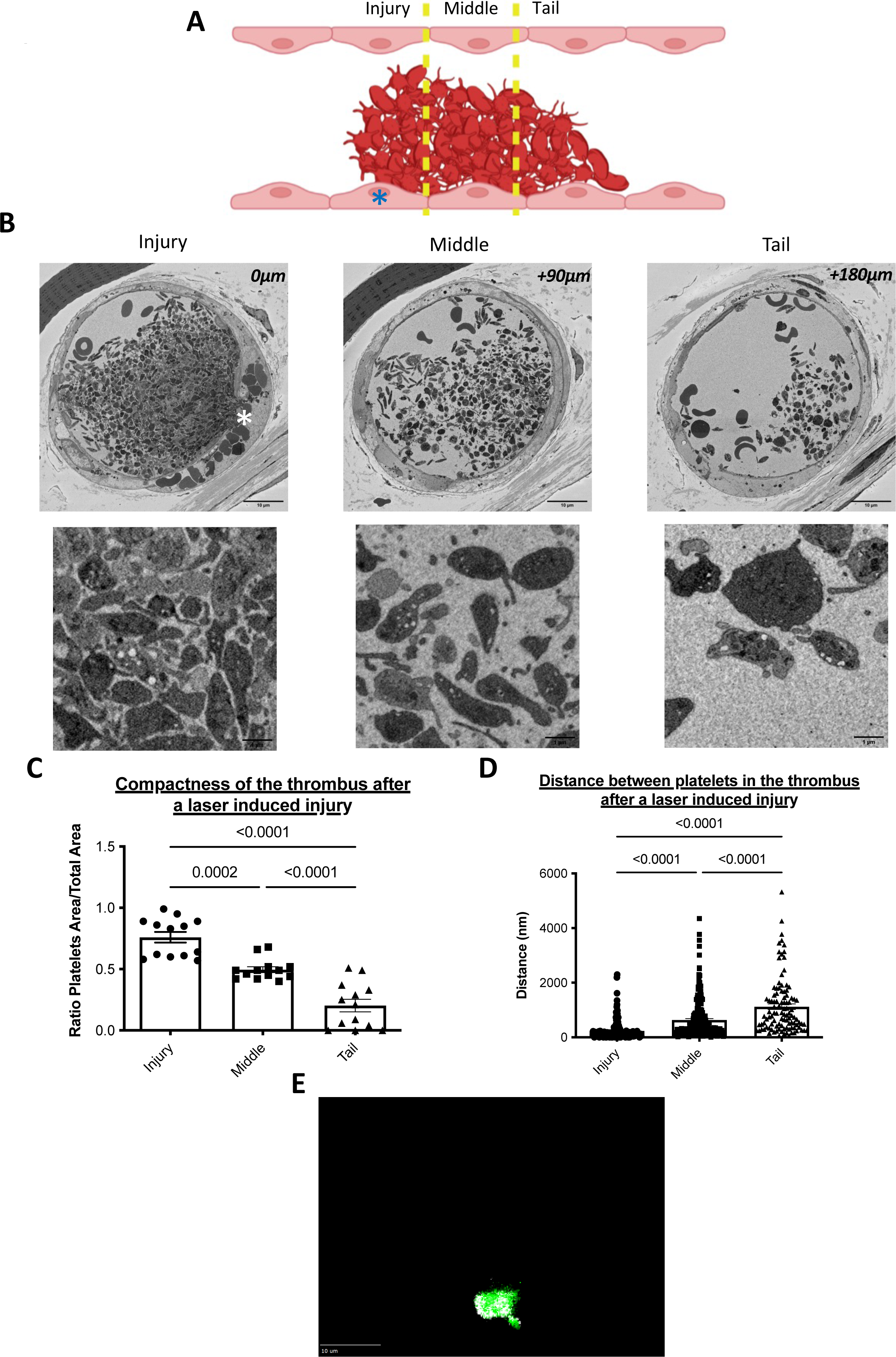
Radial gradients of platelet activation emanate from the injury site within thrombi formed after a laser-induced injury. (A): A descriptive schematic of a longitudinal thrombus section obtained after a laser-induced injury. The site of laser injury is indicated by an asterisk. Three regions of the thrombus were defined: injury, middle, and tail. Under the diagram, representative images of a thrombus section for each region were obtained after a laser-induced injury. (B): Representative pictures of platelets by SBF-SEM enlargements for each region. (C): Quantitative analysis of thrombus compactness values in the three studied regions. 13 fields were studied in each region. The mean +/- SEM is represented. Significance was determined using one-way ANOVA with Tukey’s multiple comparisons test at 95% CI. (D): Quantitative analysis of the distances between closest neighbors’ platelets from the thrombus in the three studied regions. More than 100 distance measurements between platelets were analyzed in each region (155 in the injury, 203 in the middle, and 106 in the tail). The mean +/- SEM is shown. Significance was determined using one-way ANOVA with Tukey’s multiple comparisons test. (E) Representative projection image from 25 thrombi analyzed in three different mice of platelets (GPIbβ, in green) and TLT-1 (in white) in thrombi following infusion of anti-GPIbβ and anti-TLT-1 antibodies.

The value corresponding to the compactness decreases from the injury (0.76), the middle (0.50) to the tail of the thrombus (0.20) (**Figure 4C**). Conversely, the distance between two neighboring platelets increases from the injury to the tail region. On average, in the injury section, there are 240 nanometers between any two platelets in the injury part, 630 nanometers in the middle, and 1100 nanometers in the tail of the thrombus (**Figure 4D**). These two parameters indicate the presence of a very dense and compact platelet network at the site of the laser injury. At the tail of the thrombus, platelets are more distant, and the thrombus is extended. These results indicate, in addition to a transversal gradient of activation, a longitudinal gradient of activation of platelets from the injury to the tail of the thrombus. Our results show that platelets are distributed in a radial fashion emanating from the thrombus core, and their activation state determines this. To confirm that activated platelets are distributed following a spatial gradient rather than a core and a shell, we next performed intravital experiments using an antibody directed against TREM-like transcript 1 (TLT-1) to detect activated platelets without using Fura-2. We observed, as previously published [17], that TLT-1 is more rapidly translocated to the surface of activated platelets than P-selectin during thrombus formation in vivo (data not shown). In addition, using this antibody, activated platelets (in white) were detected both in the core of the thrombus but also at the periphery (**Figure 4E**), confirming our results obtained with the calcium flux (Fura-2) and analysis by computational biology.

### Fibrin is generated independently of the presence and activation of platelets at the site of the injured endothelium.

Our results indicate that most platelets at the injury site (in the first or core region) are activated and form filopodia. Fibrin is mainly generated at this region [18][19], suggesting that platelets present in the core of the thrombus may be procoagulant. Previous results demonstrated that fibrin generation is not affected in PAR4 null mice in the Furie Model [20]. Using LSCIM, we observed that fibrin is mainly colocalized with the injured endothelium and not platelets (**Figure 5A**). Only 6% of the fluorescent signal corresponding to platelets co-localized with the fluorescent signal corresponding to fibrin, versus 70% of colocalization for fibrin and the injured endothelium (**Figure 5B**), suggesting that platelets are not involved in fibrin generation in this model of thromboinflammation. We next used the R300 antibody in the conditions in which we previously demonstrated than more than 95% of circulating platelets were depleted following its infusion in the bloodstream but not following the infusion of the isotype control [19]. We compared the kinetics of fibrin generation in P2Y12 KO mice (in which platelet activation is strongly inhibited [18][21]) and WT control (**Figure 5C**). As expected, platelet accumulation was strongly and significantly reduced in P2Y12 KO mice compared to control mice (**Figure 5D top panel**). However, fibrin generation was unchanged (**Figure 5D bottom panel**), indicating that the presence and activation of platelets is not required for fibrin generation in this model of thrombosis.

**Figure 5:**
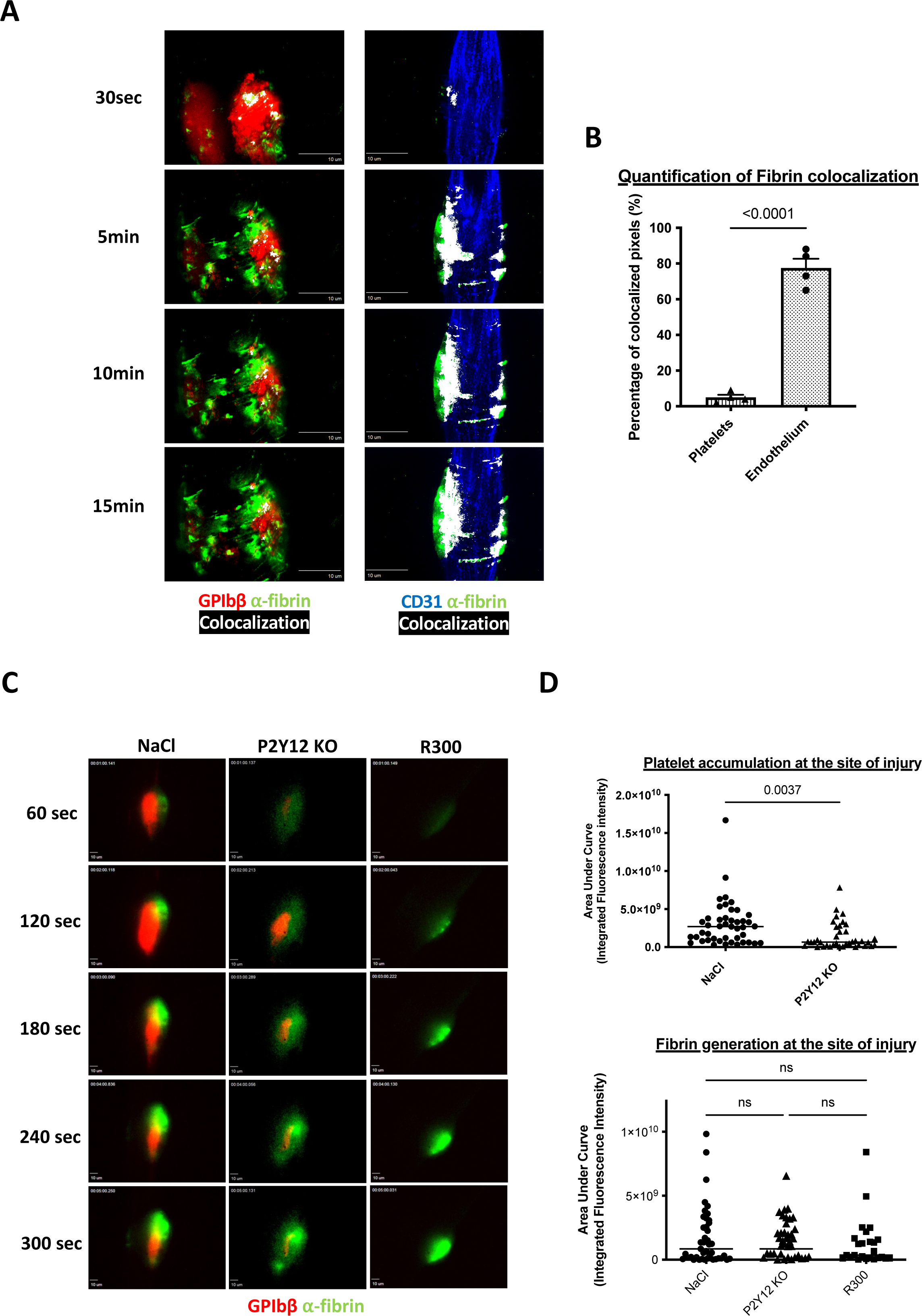
Platelets are not necessary for the activation of the coagulation cascade and the generation of fibrin *in vivo*. (A): Representative images of the colocalization between platelets and fibrin (left panel) and fibrin and endothelium (right panel) in LSCIM after laser-induced injury over time in wild-type mice. Left and right panels are independent experiments. Left panel: platelets (in red) are revealed by an antibody against the GPIbβ protein directly coupled to DyLight 649 (0.25 μg/g of the mouse) and the fibrin (in green) is visualized by an antibody directed against the fibrin labeled with a DyLight 488 (0.25 μg/g of the mouse). Right panel: the endothelium (in blue) is detected using an antibody against CD31 coupled to an AlexaFluor 647 (0.25 μg/g of the mouse) and the fibrin (in green) is visualized by an antibody directed against the fibrin labeled with a DyLight 488 (0.25 μg/g of the mouse). Antibodies are injected in IV before performing the laser-induced injury and the thrombi are followed every 3 minutes for a total duration of 15 minutes. The colocalization of the two partners (platelet/fibrin or endothelium/fibrin) is shown in white. Scale bars: 10μm. (B): Quantitative analysis of colocalizations between fibrin and platelets and fibrin and endothelium. The graph represents the mean +/- SEM of the percentages of colocalized pixels under these two conditions (fibrin/platelets or fibrin/endothelium at the final time point 15min). 4 mice per condition were analyzed and the significance was determined using an unpaired two-tailed Student’s t-test. (C): Representative images of the kinetics of fibrin generation by intravital microscopy in wild-type mice in the presence of platelets (NaCl, left panel), in mice with platelets that do not express P2Y12 receptor (ADP receptor) (P2Y12 KO, middle panel) and wild-type mice depleted in platelets with the antibody anti-GP1bα (R300, right panel). Fibrin represented in green was labeled with an anti-fibrin antibody coupled to a DyLight 488 (0.25 μg/g of the mouse). Platelets were labeled with an anti-GPIbβ antibody coupled to a DyLight 649 and administered in an amount of 0.25 μg/g of the mouse. Scale bars: 10μm. (D): Statistical analysis of the areas under the curve of integrated fluorescence intensities corresponding to the platelet accumulation (top panel) and fibrin generation (bottom panel). Data are shown as medians, and statistics were performed using a one-way ANOVA with Mann-whitney test at 95% CI. (NaCl: 45 thrombi in four mice; P2Y12 KO: 38 thrombi in three mice; R300: 25 thrombi in three mice).

To identify the presence of a procoagulant and negatively charged phospholipid surface at the site of injury, we infused Annexin-V in mice before carrying out a laser-induced injury with a confocal intravital microscope (**Figure 6**). Following the laser-induced injury, the fluorescent signal corresponding to the presence of negative phospholipids was detected at the injury site and colocalized with the endothelium (**Figure 6A top panel).** The depletion of circulating platelets using R300 antibody, did not affect Annexin-V accumulation on the injured endothelium (**Figures 6A bottom panel and 6B**). We concluded that following a laser-induced injury, the endothelium, but not platelets, constitute the catalytic surface necessary for activating the blood coagulation cascade leading to thrombosis.

**Figure 6:**
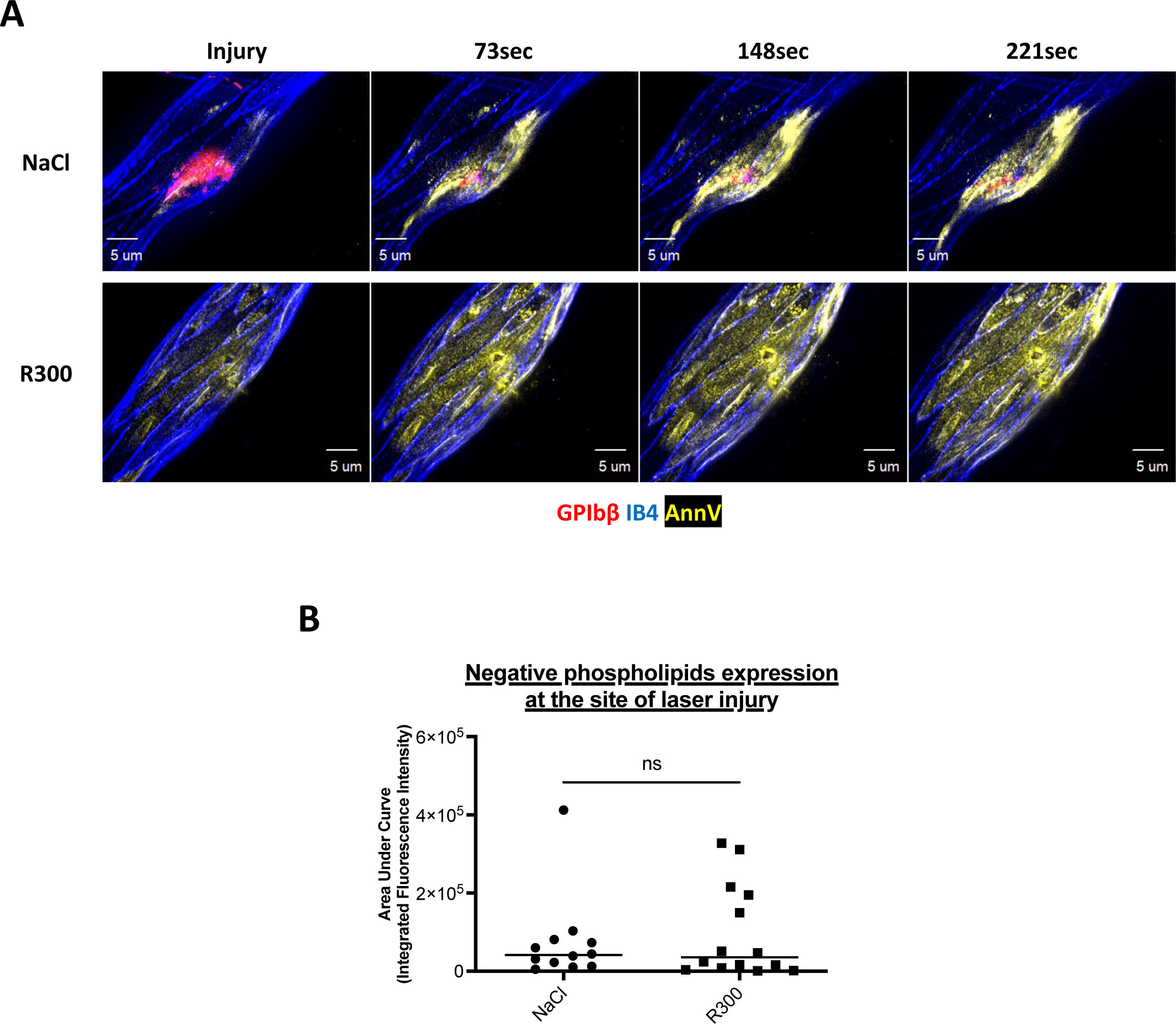
The activated endothelium is the catalytic surface rich in negative phospholipids on which the activation of the coagulation cascade occurs. (A): Representative images of the expression of negative phospholipids by LSCIM after a laser-induced injury over time in wild-type mice in the presence of platelets (NaCl, top panel) and the absence of platelets, deleted with R300 antibody (R300, bottom panel). Platelets (in red) are localized using an antibody against the GPIbβ protein directly coupled to DyLight 649 (0.25 μg/g of the mouse). The endothelium (in blue) is detected using Isolectine-GS IB4 directly conjugated to an Alexa Fluor 568 (0.25 μg/g of the mouse). Phosphatidylserines (in yellow) are visualized using fluorescent Annexin-V directly labeled with Pe-Cy5. Antibodies are injected in IV through the jugular vein before performing the laser-induced injuries and the thrombi are followed every 70 seconds over several minutes. Scale bars: 5μm. (B): Statistical analysis of the areas under the curve of integrated fluorescence intensities over times corresponding to the expression of negative phospholipids. Data are shown as median, and the significance was determined using a one-way ANOVA with Mann-whitney test (95% of confidence. (NaCl: 12 thrombi in three mice; R300: 14 thrombi in three mice).

Altogether, our results show that platelets are distributed in a radial fashion emanating from the thrombus core, and their activation state determines this. However, resting platelets could be present at the injury site, like fully activated platelets may be present in the periphery area. The state of platelet activation correlates with the duration of a platelet in a thrombus. Finally, in this model of thromboinflammation, the injured endothelium but not platelets constitute the primary source of negative phospholipids needed to activate the TF-dependent coagulation cascade.

## Discussion

The concept of immunothrombosis was coined ten years ago when our team, using the Furie model [19], and the Massberg team, using the DVT model [22][23][24], showed that granulocytes and in particular neutrophils play a role in thrombus formation in models in which collagen is not exposed to the bloodstream. Since then, different studies have confirmed the critical role played by neutrophils (and monocytes) in thrombosis [24][25][26]. Neutrophils Extracellular Traps (NETs) in the DVT model were also described to activate both platelets and the blood coagulation cascade. However, our previous results demonstrate that DNAse-I also has ATPase and ADPase activities resulting in the inhibition of platelet and neutrophil activation, which challenges this concept [27]. Here we confirmed that, although needed for the initiation of thrombus formation, neutrophils are not present in the thrombus since we could not observe them by electron microscopy. Initially, in the DVT model, NETs were detected in the red part of the thrombus [27] but not with platelets. It will be essential to determine the spatial distribution of neutrophils, NETs, and platelets in this model to determine if NETs interact with and activate platelets or, as observed in the Furie Model, are primarily involved in the generation of thrombin.

Different studies have dissected the molecular and cellular mechanisms involved in thrombus formation in the Furie model of thrombosis. The dye-based pulsed nitrogen laser activates the endothelium [28], independently of the interaction of collagen with platelets [14]. ATP-activated neutrophils [29] play a role in this model, leading to the TF dependent – and Factor XII independent-generation of thrombin [19]. We recently confirmed the presence of the endothelium and activated neutrophils (but no neutrophil extracellular traps) at the injury site using electron microscopy [30]. When investigating the activation state of platelets participating in a growing thrombus, we previously described that thrombin is a key agonist involved in intracellular calcium mobilization in platelets [13][14]. Considering the condensation of platelets and their colocalization with P-selectin and fibrin, Stalker *et al*. [9] proposed a model in which the thrombus is divided into two distinct regions: one close to the wound, called the “core” with highly condensed, activated and procoagulant platelets and a region in the periphery called the “shell” composed of less activated (P-selectin negative) platelets. This hypothesis was rechallenged in 2019 in a model of hemostatic plug formation within the large veins [29]. The authors proposed that thrombin generated at the injury site remains in the core and only activates platelets in this region. In this study, we focused on the analysis of platelet activation in thrombus formation following an injury induced by a dye laser using computational biology (the "Furie" model of thrombosis [30]). We observed a rapid expression of negative phospholipids at the surface of the injured endothelium. Platelets accumulating at the site of thrombosis may be resting, partially activated, forming pseudopods but not procoagulant. The degree of platelet activation correlates with the time a platelet spends in the growing thrombus. We concluded that rather than two distinctive regions, a core and a shell, a thrombus is composed of a dynamic gradient of activated platelets from the injured endothelium to the vessel’s lumen.

Activated platelets released ADP and TxA2. These small secondary agonists could diffuse through the core, thus allowing the recruitment of new platelets into the shell. Conversely, we observed that ATP, ADP, and adenosine played a vital role in the activation of neutrophils and platelets at the site of injury – in the core. Kinetics of neutrophil and platelet accumulation were drastically reduced in mice deficient for P2X1 [31], P2Y12 [18], and when apyrase was infused in the bloodstream of a wild-type mice [32]. Here, by computational analysis quantifying the sphericity values of platelets and with 3-dimensional analysis of Fura-2 AM loaded platelets, “fully” activated platelets were also detected in the periphery – the shell – of the thrombus. Taken together, our results support a model in which the architecture of a thrombus and its composition respond to a complex and dynamic organization that appears to be organized in a radial gradient of activation instead of in two distinct sections, such as a core and shell.

Several research groups are focusing on the cellular composition of thrombi. A better understanding of the histopathological characteristics of thrombi is essential to lead to future advancements in cardiovascular disease treatment. The dynamic nature of thrombus composition changes over time, increasing the complexity of answering this question. Indeed, the local environment is the main contributor to any thrombus. Hathcock [31] assessed that local hemodynamic forces regulate thrombus formation pathways, such as blood flow, shear stress, and turbulence. Platelet-rich thrombi are usually associated with arterial beds (high-shear stress), whereas venous thrombi, formed at low-shear stress, are platelet-poor but fibrin- and red blood cells-rich. The second element to take into consideration is the thrombus age. It is now known that the life of a thrombus goes through three different stages: initiation, extension, and perpetuation. The adhesion of the platelets defines the initiation to the endothelium and the extracellular matrix: VWF expressed by the endothelium and the GPIb by the platelets are the main drivers of platelet adhesion and thrombus initiation. The extension phase is mainly made up of homophilic interactions between platelets, activated by ADP and TxA2, ATP, Gas6, serotonin, and cytokines are present in gradients. Finally, during the perpetuation, clot contraction by platelets applies forces on the fibrin network [32]. The clot contraction process forms compressed-red blood cells called polyhedrocytes [33]. These specific red blood cells regulate thrombus rigidity, permeability, and fibrinolysis resistance to the thrombus. This has been confirmed by Maly and colleagues [34] in 2022. Their study revealed the composition of thrombi aspirated from myocardial infarction patients depending on the age of the thrombi. Here we observed different morphology of platelets participating in a growing thrombus. It will be interesting to consider the phenomenon of contraction in the morphological analysis of platelets.

Numerous studies have identified platelets as an essential cellular partner for the activation of the coagulation cascade, ultimately leading to the generation of fibrin. This was mainly the result of *in vitro* aggregation experiments where, in the presence of potent agonists such as collagen and thrombin, platelets rapidly aggregate, express negative phospholipids at their surface, and form a fibrin-rich thrombus[35]. These procoagulant platelets were coined "COATed " platelets in reference to the agonists that induce them *in vitro* and were more recently studied *in vivo* in different animal models. Procoagulant platelets were also observed by inducing the exposure of collagen to circulating blood *in vivo* [36]. These procoagulant platelets were localized at the surface of thrombi caused by the topical application of ferric chloride. More precisely, Nechupurenko et al. [37] described that procoagulant platelets, initially formed at the injury site, could move during thrombus formation and end up at the periphery in a process named thrombus contraction. Then, in contact with circulating thrombin, procoagulant platelets will promote the generation of fibrin and ensure the stability of the residual thrombus.

The Furie model does not expose the subendothelial matrix to blood circulation. We now confirm that fibrin does not co-localize with the platelet-rich thrombus, and that fibrin generation is unaffected by a lack of platelets. Our results indicate that platelets are not involved in activating the blood coagulation cascade. These results agree with what was observed by Dr. Vivien Chen’s team in 2015 [36], who studied the role of platelets in the activation of the blood coagulation cascade when thrombosis is induced independently of subendothelial matrix exposure. Additionally, in the Furie model, Ivanciu et al. have previously shown that prothrombinase is distributed away from platelets and is largely found on activated endothelium [38]. Therefore, based on these results, we hypothesize that the activated endothelium and not platelets play a vital role in fibrin generation and the activation of the blood coagulation cascade. Although it might not be surprising that, in the absence of collagen exposure, platelets are not procoagulant, it is important to consider that fibrin could be formed independently of platelets. This may be relevant in the case of immunothrombosis and thromboinflammation.

Altogether our results indicate that a thrombus is a dynamic structure formed by resting, partially, and fully activated platelets. The degree of activation correlates with the participation of the platelets in the thrombus. Fibrin may develop independently of this activation gradient. Comparing these results with other thrombosis models will be essential to better understand the dynamics of platelet activation in a growing thrombus.

## Acknowledgments

The authors deeply acknowledge Barbara and Bruce Furie for their mentorship, constant support, and scientific advice while preparing the manuscript. The authors gratefully acknowledge Perrine Chaurand and Daniel Borschneck (CEREGE, Aix-Marseille University) for their help to localize thrombi in mouse cremaster by X-ray microcomputed tomography. The electron microscopy experiments were performed on the PiCSL-FBI core facility (IBDM, AMU-Marseille), member of the France-BioImaging national research infrastructure (ANR-10-INBS-0004). We would also like to thank Nathan Asquith and Roelof Bekendam for their critical reading and comments on the manuscript.

## Author Contribution Statements

E.C. designed the research, performed experiments, analyzed the data, and drafted and critically edited the manuscript. J.T, L.C. and N.B. performed experiments and critically edited the manuscript. G.M.S. and A.M. designed the research and critically edited the manuscript. C.D. and L.P.D. acquired the funding, designed the research, and drafted and critically edited the manuscript.

---

**Supp Fig 1.**
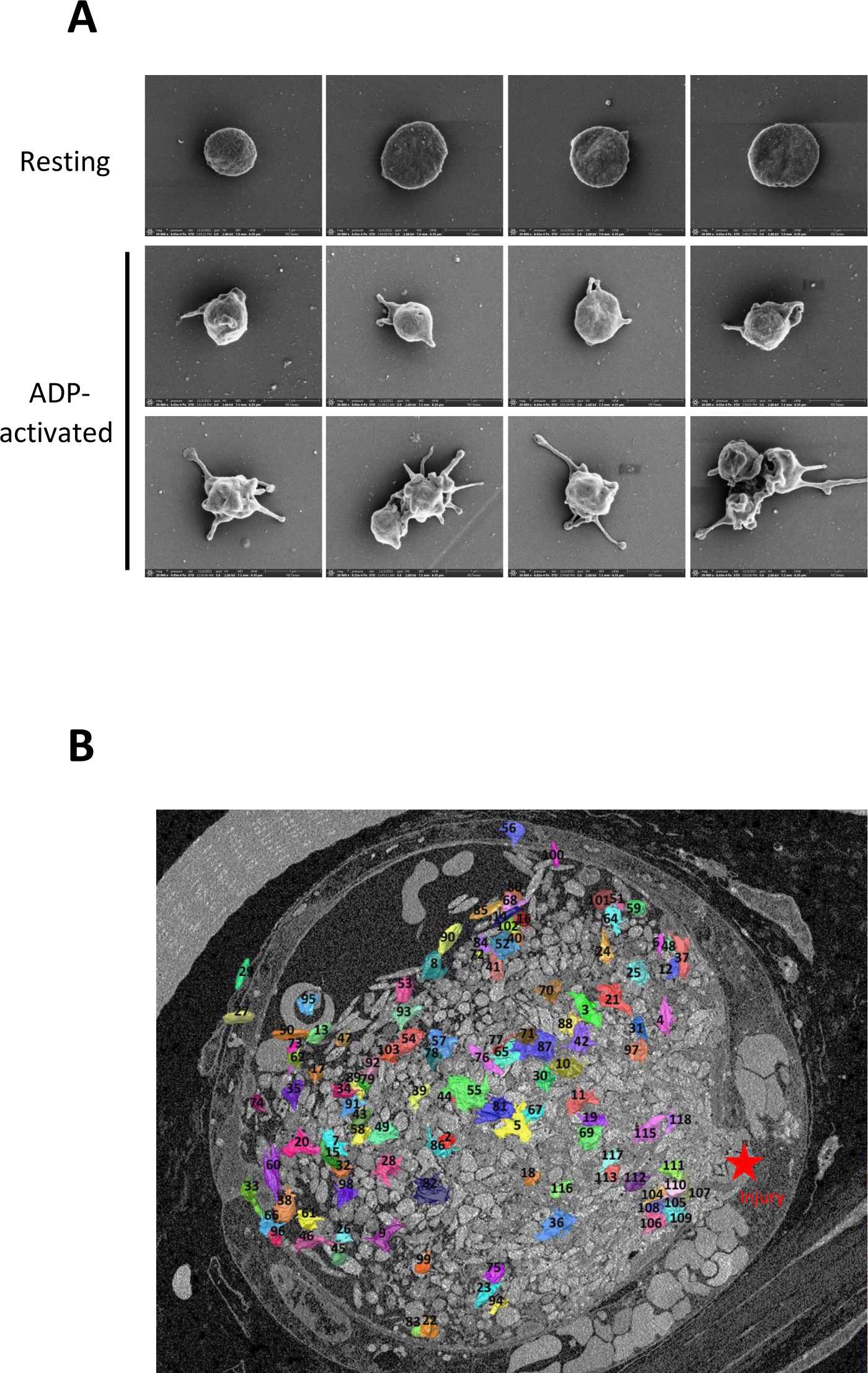

